# Systematic characterization and genetic interaction analysis of adhesins in *Candida albicans* virulence

**DOI:** 10.1101/2020.10.22.350991

**Authors:** Sierra Rosiana, Liyang Zhang, Grace H. Kim, Alexey V. Revtovich, Arjun Sukumaran, Jennifer Geddes-McAlister, Natalia V. Kirienko, Rebecca S. Shapiro

## Abstract

*Candida albicans* is a microbial fungus that exists as a commensal member of the human microbiome and an opportunistic pathogen. Cell surface-associated adhesin proteins play a crucial role in *C. albicans’* ability to undergo cellular morphogenesis, develop robust biofilms, colonize, and cause infection in a host. However, a comprehensive analysis of the role and relationships between these adhesins has not been explored. We previously established a CRISPR-based platform for efficient generation of single- and double-gene deletions in *C. albicans*, which was used to construct a library of 144 mutants, comprising 12 unique adhesin genes deleted singly, or in every possible combination of double deletions. Here, we exploit this adhesin mutant library to explore the role of adhesin proteins in *C. albicans* virulence. We perform a comprehensive, high-throughput screen of this library, using *Caenorhabditis elegans* as a simplified model host system, which identified mutants critical for virulence and significant genetic interactions. We perform follow-up analysis to assess the ability of high- and low-virulence strains to undergo cellular morphogenesis and form biofilms *in vitro*, as well as to colonize the *C. elegans* host. We further perform genetic interaction analysis to identify novel significant negative genetic interactions between adhesin mutants, whereby combinatorial perturbation of these genes significantly impairs virulence, more than expected based on virulence of the single mutant constituent strains. Together, this yields important new insight into the role of adhesins, singly and in combinations, in mediating diverse facets of virulence of this critical fungal pathogen.

**Summary:** *Candida albicans* is a human fungal pathogen and cause of life-threatening systemic infections. Cell surface-associated adhesins play a central role in this pathogen’s ability to establish infection. Here, we provide a comprehensive analysis of adhesin factors, and their role in fungal virulence. Exploiting a high-throughput workflow, we screened an adhesin mutant library using *C. elegans* as a simple model host, and identified mutants and genetic interactions involved in virulence. We found that adhesin mutants are impaired in *in vitro* pathogenicity, irrespective of their virulence. Together, this work provides new insight into the role of adhesin factors in mediating fungal virulence.

## Introduction

Fungal pathogens have been emerging as a significant threat to human health, resulting in over 1.6 million deaths worldwide each year (Denning 2017; Bongomin *et al.* 2017; Geddes-McAlister and Shapiro 2019). Despite fungal pathogens affecting over 1 billion people each year, they still remain relatively understudied compared with many other infectious disease pathogens (Denning 2017; Fisher *et al.* 2020). *Candida albicans* is amongst the most pervasive fungal pathogens of humans, and can cause infectious disease ranging from acute mucosal infections, to systemic candidiasis with extremely high morbidity and mortality rates (Pfaller and Diekema 2007; Kullberg and Arendrup 2016; Bongomin *et al.* 2017). *C. albicans* is an opportunistic pathogen present in the gastrointestinal tract, skin, reproductive tract, and oral cavity of most healthy adults. It asymptomatically colonizes many tissues of the human body, and may overgrow if there is a perturbation or depression of the host immune system, including treatment with antibiotics, organ transplants in combination with immunosuppressive drugs, or diseases such as HIV/AIDS (Kullberg and Arendrup 2016).

The success of *C. albicans* as a human pathogen relies on multiple virulence strategies, including morphological plasticity and robust biofilm formation (Shapiro *et al.* 2011; Sudbery 2011; Mayer *et al.* 2013). *C. albicans* is a polymorphic yeast, and its ability to reversibly transition between yeast and filamentous growth states (including hyphal and pseudohyphal growth) is a critical component of this pathogen’s virulence (Sudbery 2011). *C. albicans* is capable of forming robust biofilms, not only on host tissues, but on hospital equipment and medical implants such as catheters, pacemakers, and prosthetics (Finkel and Mitchell 2011; Nobile and Johnson 2015; Lohse *et al.* 2018). With the rising usage of medical implants, instances of implant-related infections are on the rise, with the majority of these infections associated with microbial biofilms (Finkel and Mitchell 2011; Nobile and Johnson 2015; Tsui *et al.* 2016; Lohse *et al.* 2018). As with many other microbial pathogens, *C. albicans* biofilms are typically resilient to many external stressors such as antifungals and host defense factors, making *C. albicans* significantly more difficult to treat in a biofilm state (Nobile and Johnson 2015; Tsui *et al.* 2016; Sharma *et al.* 2019).

*C. albicans* pathogenesis is significantly impacted by its adherence abilities, and indeed the most frequently isolated pathogenic *Candida* species are those with the greatest adhesive capacities, and these tend to be more pathogenic than other strains (Calderone and Braun 1991; Hoyer 2001). While many factors are involved in *C. albicans* adhesion, it is known that this process is largely due to the expression of fungal cell wall proteins, including adhesins, which are highly-expressed on filamentous cells, and involved in surface adhesion, biofilm formation, and host colonization (Sundstrom 1999; de Groot *et al.* 2013a; Lipke 2018). A family of adhesin proteins of particular interest in *C. albicans* is the *ALS* (agglutinin-like sequence) family of cell surface glycoproteins (Hoyer 2001). This family shares a three domain structure consisting of a high-complexity N-terminal domain that mediates protein-ligand interactions with host cells or other substrates, a low-complexity central domain that is highly variable in length, and a C-terminal domain that anchors the adhesin to the fungal cell wall via a glycosylphosphatidylinositol (GPI) anchor (Hoyer 2001; Hoyer and Cota 2016). There are eight *ALS* loci currently described in *C. albicans*: *ALS1-7* and *ALS9* (Hoyer and Cota 2016). Other families of adhesins have also been identified, including *HWP*, *IFF* and *HYR* (de Groot *et al.* 2013b). Some *C. albicans* adhesins, such as *ALS1* and *ALS3*, have been subject to fairly comprehensive molecular genetic and biochemical analysis, and have well-described roles in various aspects of adhesion, host-pathogen interactions, filamentation, and fungal virulence (Fu *et al.* 1998, 2002; Hoyer *et al.* 1998; Zhao *et al.* 2004; Sheppard *et al.* 2004; Ibrahim *et al.* 2005; Phan *et al.* 2007; Nobile *et al.* 2008; Almeida *et al.* 2008; Donohue *et al.* 2011; Cleary *et al.* 2011), while other adhesins remain incompletely described or fully uncharacterized. Additional evidence indicates that the numerous *C. albicans* adhesins have complex interactions and may have complementary, compensatory, or redundant functions (Zhao *et al.* 2004, 2005; Nobile *et al.* 2008; Shapiro *et al.* 2018a).

Given the complex interplay between *C. albicans* adhesin proteins, a powerful strategy to assess the function of these factors is through genetic interaction analysis. Genetic interaction analysis is a powerful strategy that typically takes advantage of single- and double-gene deletion strains to assess epistatic interactions between genes, and can be used to organize gene products into pathways, identify genetic synergies and redundancies, and predict gene function (Dixon *et al.* 2009; Baryshnikova *et al.* 2013). While genetic interaction analysis has been exploited widely in model organisms (Butland *et al.* 2008; Costanzo *et al.* 2010, 2016; Babu *et al.* 2011; Norris *et al.* 2017), it has more recently been employed as a powerful tool to dissect genetic interaction networks in diverse microbial pathogens, including *C. albicans* (Glazier *et al.* 2017, 2018; Shapiro *et al.* 2018a; b; Glazier and Krysan 2020). Our previous work established CRISPR-Cas9-based gene drive array (GDA) platform, which permits facile, precise and efficient creation of combination gene knockouts in *C. albicans,* which we applied to construct a library of 144 mutants, comprising 12 unique adhesin genes deleted singly, and in every possible combination of double deletions (Shapiro *et al.* 2018a; Halder *et al.* 2019). This library enables the analysis of complex genetic interactions between adhesins and their prospective roles in *C. albicans*’ adhesion; it further enables the identification of combinations of genes which, when deleted together, may interfere with fungal biofilm formation, host-pathogen interactions, or virulence.

Studying putative *C. albicans* virulence factors, such as adhesins, requires the use of a model host to assess fungal pathogenicity *in vivo*. *Caenorhabditis elegans* is a free-living nematode that has been exploited as a simple and practical model for studying host-pathogen interactions with diverse microbial pathogens (Aballay and Ausubel 2002; Marsh and May 2012; Issi *et al.* 2017; Kumar *et al.* 2020), including *C. albicans* (Pukkila-Worley *et al.* 2009, 2011; Jain *et al.* 2013; Elkabti *et al.* 2018; Feistel *et al.* 2019) and other fungal pathogens (Mylonakis *et al.* 2002b; Tang *et al.* 2005; Huang *et al.* 2014; Ahamefule *et al.* 2020; Hernando-Ortiz *et al.* 2020). *C. elegans* is a simple and cost-effective model organism that is readily propagated and stored, and lends itself to high-throughput screening of microbial pathogens (Moy *et al.* 2009; Kirienko *et al.* 2013, 2016). *C. elegans* host-pathogen interactions have many conserved features that are shared with mammalian species, making it an advantageous model for studying human disease and infections. The intestinal epithelial cells of *C. elegans* have morphological features, such as microvilli, that are similar to mammalian epithelial cells, and it is estimated that 40-60% of genes in *C. elegans* have human orthologs (C. elegans Sequencing Consortium 1998; McGhee 2007; Kumar *et al.* 2020). With regards to *C. albicans* infection, many biological mechanisms by which *C. albicans* infects *C. elegans* are similar in humans and nematodes, and *C. elegans* host immune responses to pathogens are remarkably conserved (Gravato-Nobre and Hodgkin 2005; Kim and Ausubel 2005; Pukkila-Worley and Mylonakis 2010). Once ingested by *C. elegans*, *C. albicans* can cause a persistent lethal infection, making monitoring the infection relatively simple for research purposes (Pukkila-Worley and Mylonakis 2010; Elkabti *et al.* 2018). Many *C. albicans* genes that are known to be required for murine infection are similarly required for infection in *C. elegans* (Pukkila-Worley *et al.* 2009; Elkabti *et al.* 2018), suggesting the utility of this model for studying fungal virulence factors.

Here, we perform systematic characterization and genetic interaction analysis of a *C. albicans* library of adhesin (or adhesin-like) gene mutants (de Groot *et al.* 2013a), deleted for these factors singly or in combinations of double knockouts. We use *C. elegans* as a model host system to perform high-throughput screening of our *C. albicans* library of 144 adhesin mutant strains, and identify single and combination genetic mutations that significantly alter fungal virulence. Following up on the strains with the highest and lowest levels of virulence, we characterize phenotypes associated with other virulence traits, including host colonization, biofilm formation, and cellular morphogenesis. We further perform genetic interaction analysis to identify and characterize significant negative genetic interactions, whereby double mutants are significantly impaired in virulence based on what we would predict given the virulence of their single mutant counterparts. We find the requirement for certain adhesins to be environmentally-contingent, and observe that many virulence-associated phenotypes are uncoupled under our experimental conditions. Together, comprehensively characterizes the roles of single- and double-adhesin mutant strains in fungal pathogenicity, with important implications for understanding fungal virulence and host interactions.

## Results

### C. elegans infection assay identifies avirulent C. albicans adhesin mutants

First, we aimed to assess the virulence profiles of a library of *C. albicans* adhesin single- and double-gene deletion mutants (Shapiro *et al.* 2018a), to assess the role of these factors, singly or in combination, in fungal pathogenicity. This adhesin mutant library consisted of 144 adhesin mutants, representing 12 single adhesin gene deletions, and 66 double adhesin gene deletions with ‘reciprocal pairs’ (*i.e. a*Δ*/b*Δ and *bΔ/aΔ*). In order to assess virulence, we used a high-throughput screening model with *C. elegans* as a model host species. Young adult *C. elegans* worms were added to 384-well plates containing each of the *C. albicans* mutant strains, and incubated for 72 hours. After infection, plates were washed to remove *C. albicans* and dead *C. elegans* were stained with a cell-impermeant fluorescent dye (**Figure 1a** and see Materials and Methods for additional details). This infection assay was repeated in six replicates. Increased survival of the *C. elegans* host indicated less virulence by the particular mutant strain of *C. albicans*. **Figure 1b** depicts an example of bright field imaging that allows visualization of all worms after infection. **Figure 1c** depicts examples of fluorescent images, where only dead *C. elegans* worms are stained and are visible with fluorescence. Worm death is monitored and calculated based on the ratio of the area of fluorescence to the area of total worms in brightfield.

**Figure 1.**
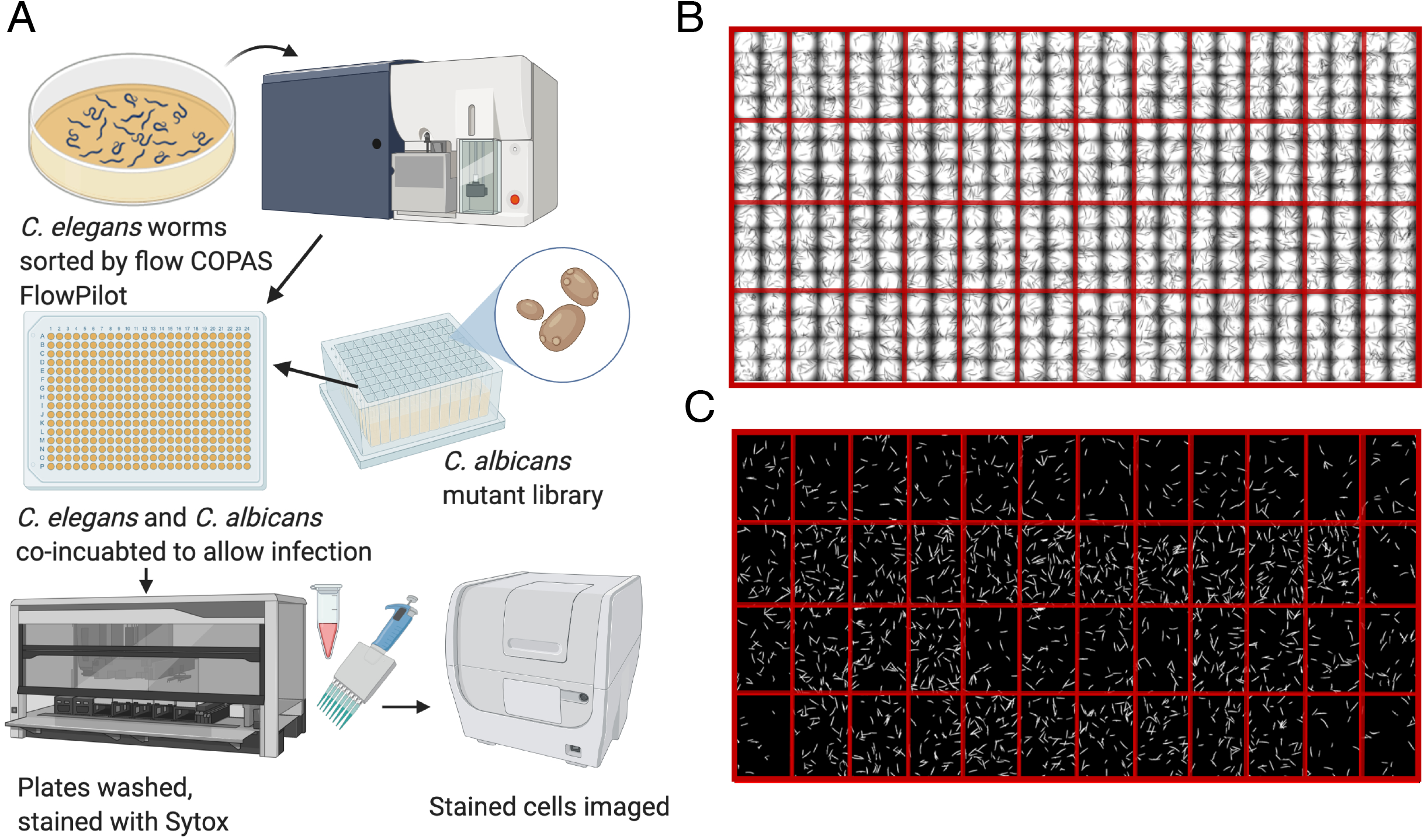
A protocol of *C. albicans - C. elegans* high-throughput screening for fungal virulence regulars. *C. elegans* is a model host for *C. albicans* fungal infection and can be used in high-throughput screening settings to identify regulators of fungal virulence. **(A)** A schematic indicating the workflow for the *C. albicans - C. elegans* infection assay. Worms were sorted by COPAS FP worm sorter and an equal number of worms were deposited into each well of a 384-well plate. *C. albicans* mutant strains (the single- and double-gene deletion library) were inoculated into the 384-well plate, and *C. albicans* and *C. elegans* were co-incubated to allow for infection. To stop the infection progression, plates were washed with a plate washer to remove *C. albicans*, and *C. elegans* were stained with a cell impermeant dye Sytox Orange to identify dead worms. Plates were then imaged in brightfield and fluorescence channels, and processed using Cell Profiler software to identify the total area of worms, as well as the area of fluorescence inside of the worms per each well. Figure was created using BioRender (biorender.com) **(B)** An example of brightfield imaging, which allows visualization of all of the worms (alive and dead) after infection. Red lines have been added to the image to depict the delineation between wells with the same strain of *C. albicans* inoculated. **(C)** Example of fluorescence microscopy images used to identify dead fluorescent worms stained with the cell-impermeant fluorescent dye, Sytox Orange. Only dead *C. elegans* worms were stained and visible with fluorescence. Red lines have been added to the image to depict the delineation between wells with the same strain of *C. albicans* inoculated (technical replicates).

The wild-type strain, with no gene deletions, was used to determine baseline virulence, and was compared with the 144 mutant strains for ability to cause death in the *C. elegans* host. As predicted, the wild-type strain showed a high level of virulence towards *C. elegans* with greater than 50% of worms killed during the course of the infection assay (**Figure 2, Table S1**). Overall, many of the adhesin mutants were found to be impaired in virulence, compared with the wild-type strain (**Figure 2**): 24 out of 78 unique genotypes (12 single mutants and 66 double mutant genotypes) led to significant reduction in worm death compared to the wild-type strain (*P* < 0.05, ANOVA, **Table S1**). Of these mutants with significantly impaired virulence, 22 were double mutants, and two (*als1Δ* and *als5Δ*) were single mutants, indicating the importance of monitoring virulence in single as well as double mutant strains. Mutation of *als1Δ*, and to a lesser extent *als5Δ*, singly or in combination with other genes, attenuated the ability of the pathogen to kill the host to the greatest extent, suggesting that these adhesin genes significantly contribute to *C. albicans*’ virulence.

**Figure 2.**
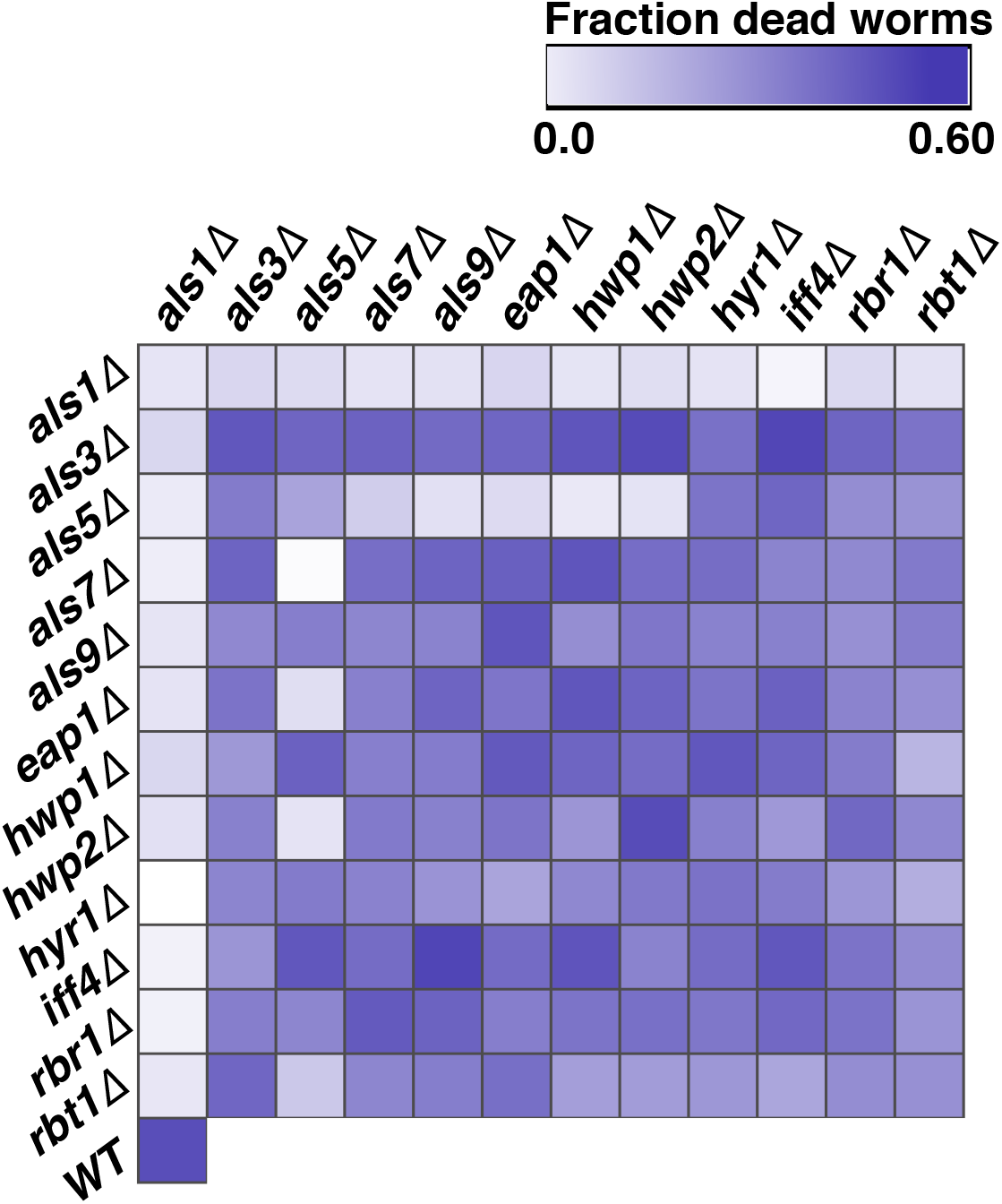
Multiple *C. albicans* adhesin mutant strains are impaired in virulence in *C. elegans* model of infection. When screened for virulence using *C. elegans* as a model host, *C. albicans* adhesin mutant strains displayed variable levels of virulence. The library of 144 single- and double-genetic mutant strains was screened for virulence, and the fraction of dead worms was established for each mutant strain. The heat map depicts the compiled and processed results from the experiment, with darker blue/purple squares indicating more worm death (higher fraction of dead worms), and white or lighter coloured squares indicating less death (lower fraction of dead worms). The heatmap shows virulence for each strain, averaged over at least four biological replicates. The heat map was generated using Morpheus matrix visualization and analysis software from the Broad Institute (https://software.broadinstitute.org/morpheus).

The five most virulent strains (excluding wild-type) were *als3Δiff4Δ*, *als9Δiff4Δ*, *hwp2Δ*, *als3Δhwp2Δ* and *als9Δeap1Δ*; all of which displayed similar death percentages to the wild-type strain (>50% of worms dead). The single mutant counterpart strains of these mutants, *hwp2Δ*, *als3Δ*, *iff4Δ*, *als9Δ*, and *eap1Δ* all show comparatively high virulence, indicating that these adhesins are not essential for virulence, at least in this infection model. The five least virulent strains were *hyr1Δals1Δ*, *als7Δals5Δ*, *als1Δiff4Δ*, *iff4Δals1Δ* and *rbr1Δals1Δ*. The three most significantly attenuated strains (*hyr1Δals1Δ*, *als7Δals5Δ*, and *als1Δiff4Δ*, <10% death) were selected for follow up study. The single mutant strains, in order from least virulent to most virulent are as follows: *als1Δ*, *als5Δ*, *rbt1Δ*, *als9Δ*, *eap1Δ*, *rbr1Δ*, *hyr1Δ*, *als7Δ*, *hwp1Δ*, *iff4Δ*, *als3Δ*, *hwp2Δ*.

The double mutant ‘reciprocal pairs’ were all validated to ensure they indicate the same results, where a reciprocal pair refers to the same mutant genotype, but generated by mating opposite mating type haploids. For example, *als1Δals3Δ* and *als3Δals1Δ* should theoretically show the same results because the same two genes were deleted, though the strains were generated independently. The majority of the mutants showed very similar data results between the reciprocal pairs, similar to what had been seen in previous analysis of this library (Shapiro *et al.* 2018a). Amongst the 66 total double deletion genotypes, 13 (*hwp1Δhwp2Δ*, *hwp1Δhyr1*, *als3Δhwp1Δ*, *als3Δhwp2Δ*, *als3Δiff4Δ*, *als5Δals9Δ*, *als5Δhwp1Δ*, *als5Δrbt1Δ*, *als7Δhwp1Δ*, *als7Δrbr1Δ*, *als9Δiff4Δ*, *als9Δrbr1Δ*, *eap1Δhyr1Δ*) had a virulence difference between their reciprocal pairs of 10 percentage points or more (*i.e.* if *aΔbΔ* resulted in 45% dead worms and *bΔaΔ* resulted in 55% dead worms).

### Low- and high-virulence C. albicans mutants exhibit different ability to colonize C. elegans

Next, we examined the relationship between *C. albicans* colonization of *C. elegans* and strain virulence. As previously described, we focused on our assembled panel of six mutant strains, three with low virulence (*als1Δiff4*, *hyr1Δals5Δ*, *als7Δals5Δ*) and three with high virulence (*als3Δhwp2Δ*, *als3Δiff4Δ*, *als9Δiff4Δ*) (**Figure 3a**; *P* < 0.0001 pooled low- and high-virulence groups, Student’s *t*-test)). Sterile, young adult *glp-4(bn2)* worms were exposed to each *C. albicans* strain in a liquid pathogenesis assay for 24 hours. Worms were then collected, washed, and lysed. Serially-diluted fungi were plated on YPD agar media and colony-forming units were counted to establish the colonization potential of these different fungal mutants. Interestingly, significant differences were observed between the levels of colonization of low-virulence compared with high-virulence subsets (**Figure 3b;** *P* < 0.05 pooled low- and high-virulence groups, Student’s *t*-test)). Strains with generally lower virulence in the *C. elegans* infection assay, demonstrated increased capacity for worm colonization (**Figure 3b**). Interestingly, this contrasts with previous observations of bacterial pathogens that colonize *C. elegans*; in those cases, colonization and virulence directly (rather than inversely) correlated (Garsin *et al.* 2003; Evans *et al.* 2008; Kirienko *et al.* 2013). One possible explanation for this difference is that *C. elegans* detects pathogenic determinants or damage afflicted by the fungus during infection and responds with increased innate immune activity that restricts fungal colonization. This suggests an interesting trend amongst these fungal mutant strains that unlinks pathogen colonization potential and virulence.

**Figure 3.**
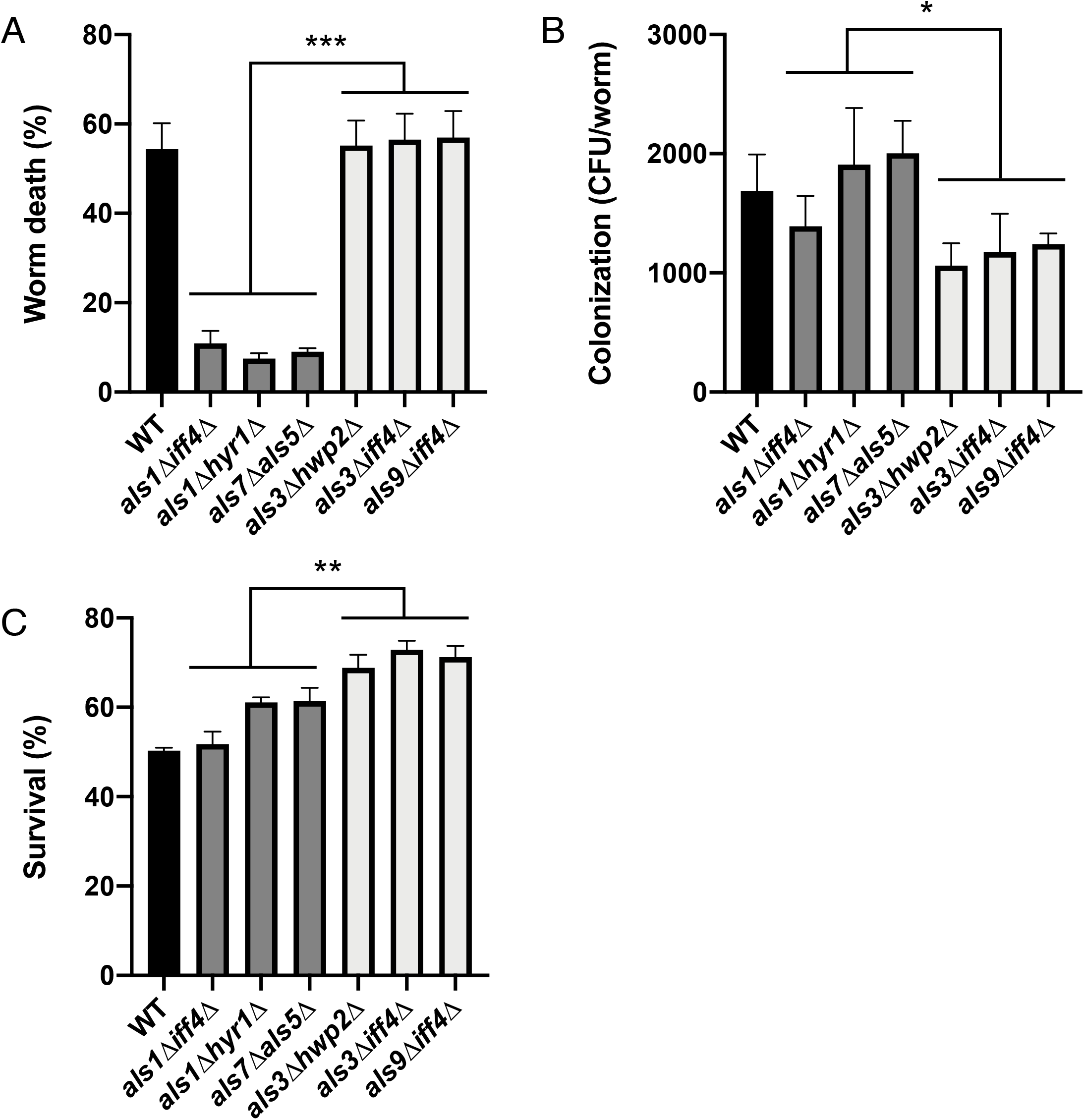
*C. albicans* mutants’ virulence depends on infection model. *C. albicans* strains with the highest and lowest levels of virulence from our *C. elegans* infection screen were selected for follow-up analysis and monitored for worm colonization, as well as virulence in an agar plate infection model. **(A)** The three strains with lowest virulence (*als1Δiff4*, *hyr1Δals5Δ*, *als7Δals5Δ*; dark grey) and three with highest virulence (*als3Δhwp2Δ*, *als3Δiff4Δ*, *als9Δiff4Δ*; light grey), based on the high-throughput *C. elegans* liquid media screen. Graph depicts the percent of worm death. The three strains with lowest virulence were significantly different from the wild-type strains *P* < 0.0001 for the pooled low- and high-virulence groups based on Student’s *t*-test (***)). **(B)** Ability of *C. albicans* mutant strains and wild type to colonize *C. elegans* worms was monitored. Colonization was assessed using colony-forming units (CFU) of *C. albicans* per worm. Lower virulence mutants (dark grey) had higher CFU/worm compared with higher virulence mutants (light grey) (*P* < 0.05 for the pooled low- and high-virulence groups based on Student’s *t*-test (*)). **(C)** An agar-based *C. elegans* infection model was used to compare data from the liquid-based infection screen. In this agar assay, *C. elegans* survival was scored daily until all the worms were dead. Graph depicts *C. elegans* survival data at day 4, at which point wild-type *C. albicans* has killed approximately 50% of worms. Low- and high-virulence mutants show a reverse trend from liquid assay, as strains that were highly virulent liquid killing had low pathogenesis on solid media, and vice versa (p<0.001 for the pooled low-and high-virulence groups based on Student’s *t*-test (**)). All graphs were generated with GraphPad Prism. Error bars represent SEM.

### Candida virulence mechanisms differ between liquid- and agar-based assays

Research using *C. elegans*-based pathogenesis models have convincingly demonstrated that the microbial virulence determinants are strongly influenced by the context of the infection assay (e.g., media composition, state of matter, etc.). For example, at least five distinct *C. elegans*–*P. aeruginosa* pathogenesis models have been described (Mahajan-Miklos *et al.* 1999; Tan *et al.* 1999; Gallagher and Manoil 2001; Zaborin *et al.* 2009; Kirienko *et al.* 2013; Utari and Quax 2013). To investigate whether this phenomenon holds true for *C. albicans*, host killing in an agar-based infection model was compared to data from the liquid-based *C. elegans* infection model. In this agar-based assay, *C. elegans* survival was scored daily until all the worms were dead. For a more direct comparison for liquid-based assay, we extracted survival data for day 4. At this time, wild-type *C. albicans* has killed approximately 50% of worms, a value consistent with our observations from liquid-based assays. Interestingly, when pooled data for low- and high-virulence mutants were compared, we saw the reverse of the outcome from the liquid-based assay: strains that were highly pathogenic in liquid killing and had low colonization, also had low pathogenesis on solid media (**Figure 3c**; *P* < 0.001 pooled low- and high-virulence groups, Student’s *t*-test)). Kaplan-Mayer curves demonstrating longitudinal survival are shown in **Figure S1**.

We hypothesized that the relative expression of adhesin genes, and perhaps the compensatory upregulation of different adhesin genes in different mutant strain backgrounds under variable growth conditions, may underlie these differences. To test this prediction, qRT-PCR was used to measure levels of four adhesin genes (*ALS1*, *EAP1*, *HYR1*, and *IFF4*) from low virulence (*als7Δals5Δ*) and high virulence (*als3Δhwp2Δ*) mutants. In addition to their phenotypes associated with virulence in the *C. elegans* liquid infection assay, these two mutants were further selected based on the highest (*als7Δals5Δ*, 2002 CFU/worm) and the lowest (*als3Δhwp2Δ*, 1060 CFU/worm) ability to colonize *C. elegans,* respectively. Expression levels were measured for each strain using both liquid and agar-based conditions.

Interestingly, of the adhesins measured, *ALS1* expression was markedly higher (under all conditions tested) than the other three genes (**Figure 4a**). This outcome was surprising, as this gene has been deleted from two of the mutants that retain high colonization ability (*hyr1Δals1Δ* and *iff4Δals1Δ*). This suggests that either this adhesin does not play a prominent role in colonization, or one or more other genes compensate for its loss, making this gene redundant. Next, we compared expression of all four adhesin genes from *als3Δhwp2Δ* and *als7Δals5Δ* mutants under liquid and agar growth conditions. In both cases, expression of *ALS1* was somewhat lower in the high colonization strain (*als7Δals5Δ*; **Figures 4b and 4c**), which is consistent with our hypothesis that *ALS1* may be dispensable for host colonization. Expression of the other three genes, *EAP1*, *HYR1*, and *IFF4*, was higher in the high colonization strain (*als7Δals5Δ*), compared to the low colonization strain (*als3Δhwp2Δ*, **Figure 4b and 4c**), though these differences were much more pronounced in liquid compared with on solid media. Taken together, these data suggest that distinct adhesin gene expression patterns are exhibited by *C. albicans* mutants with different levels of colonization, as well as different levels of virulence. In addition, our data also support the conclusion that, like bacterial pathogenesis assays, microbial physiology (as determined by growth media) can have a profound impact on pathogen virulence.

**Figure 4.**
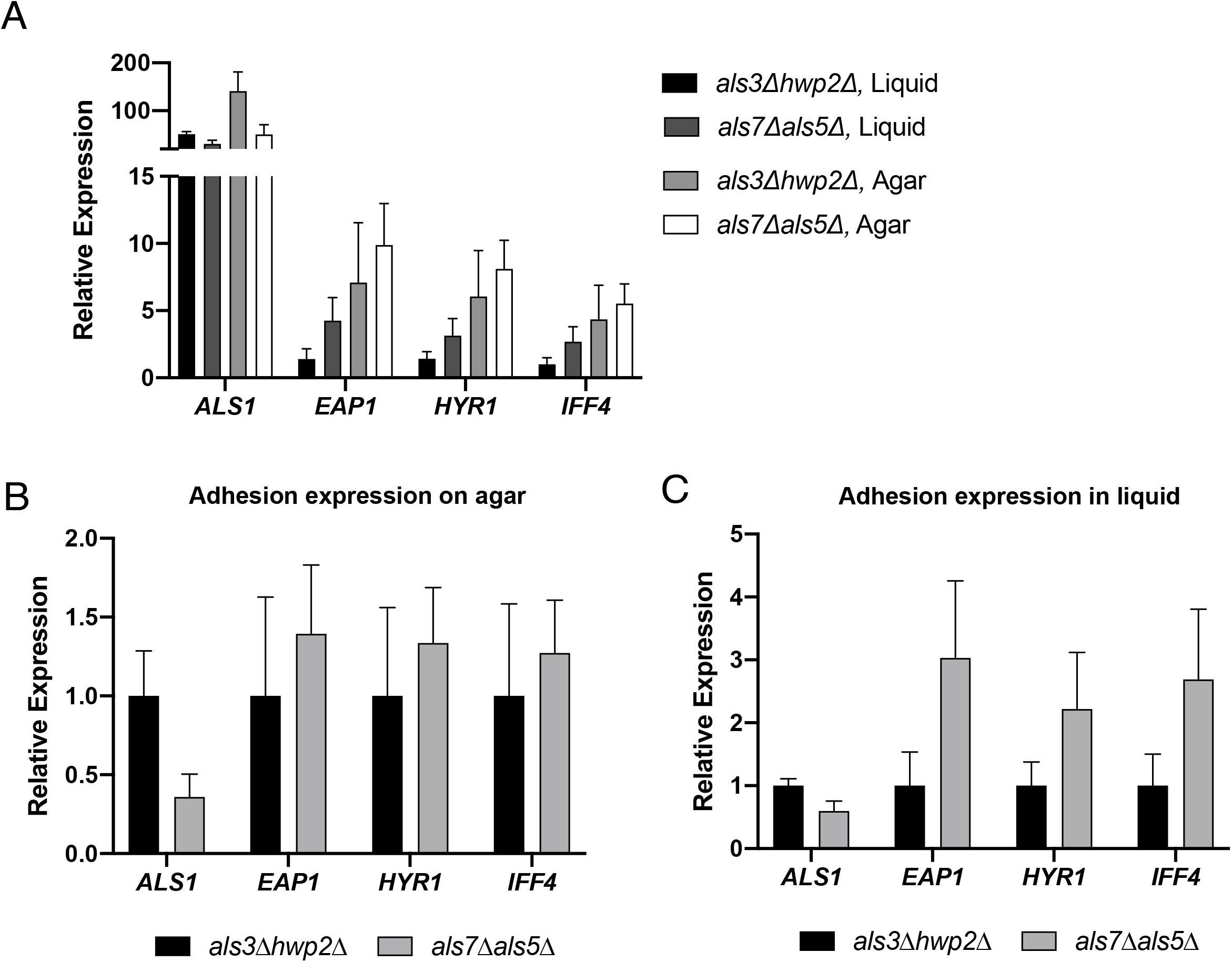
Distinct adhesin gene expression patterns are exhibited by *C. albicans* mutants with different levels of virulence and colonization. qRT-PCR was used to measure levels of four adhesin genes (*ALS1*, *EAP1*, *HYR1*, and *IFF4*) for representative low virulence, high colonization (*als7Δals5Δ*) and high virulence, low colonization (*als3Δhwp2Δ*) mutants. Expression levels were measured for each strain under both liquid and agar conditions. **(A)** Relative expression of *ALS1*, *EAP1*, *HYR1*, and *IFF4* (measured relative to the housekeeping gene *ACT1*) in high colonization / low virulence (*als7Δals5Δ*) and high virulence / low colonization (*als3Δhwp2Δ*) mutants, under liquid or solid growth conditions. Relative gene expression was further normalized to the lowest expressed gene (*IFF4* in *als3Δhwp2Δ* in liquid media). **(B-C)** Relative adhesin expression on agar **(B)** or in liquid **(C)**. Expression values in high colonizer *als7Δals5Δ* were normalized to low colonizer *als3Δhwp2Δ* for each gene. Error bars represent SEM.

### Adhesin mutant strains are impaired in filamentation and biofilm formation

Following *in vivo* analysis of fungal virulence and colonization, we performed *in vitro* biofilm and filamentation assays to determine whether observed differences in fungal virulence and/or colonization could be ascribed to altered ability of adhesin mutant strains to form biofilm or undergo cellular morphogenesis. Therefore, we first performed *in vitro* biofilm growth assays of the wild-type strain, along with the three most virulent, and three least virulent adhesin mutant strains. The selected strains were allowed to form biofilms in flat bottom 96-well plates for 72 hours, planktonic cells were removed, and metabolic activity of the remaining biofilm was measured via XTT and quantified by spectrophotometer readout at OD_490_. With the exception of *als9Δiff4Δ*, each of the adhesin mutant strains were found to have impaired biofilm formation compared with the wild-type strain (**Figure 5a;** *P* < 0.0005). Given the known role of adhesins in fungal adhesion and biofilm initiation and growth, it is expected that these mutant strains would likely be defective in biofilm growth. However these results do not indicate a correlation between biofilm formation and virulence in our *C. elegans* liquid killing or agar-based assays, and suggests that mutants with reduced ability to form biofilms under these *in vitro* conditions are still capable of virulence in the *C. elegans* model of infection.

**Figure 5.**
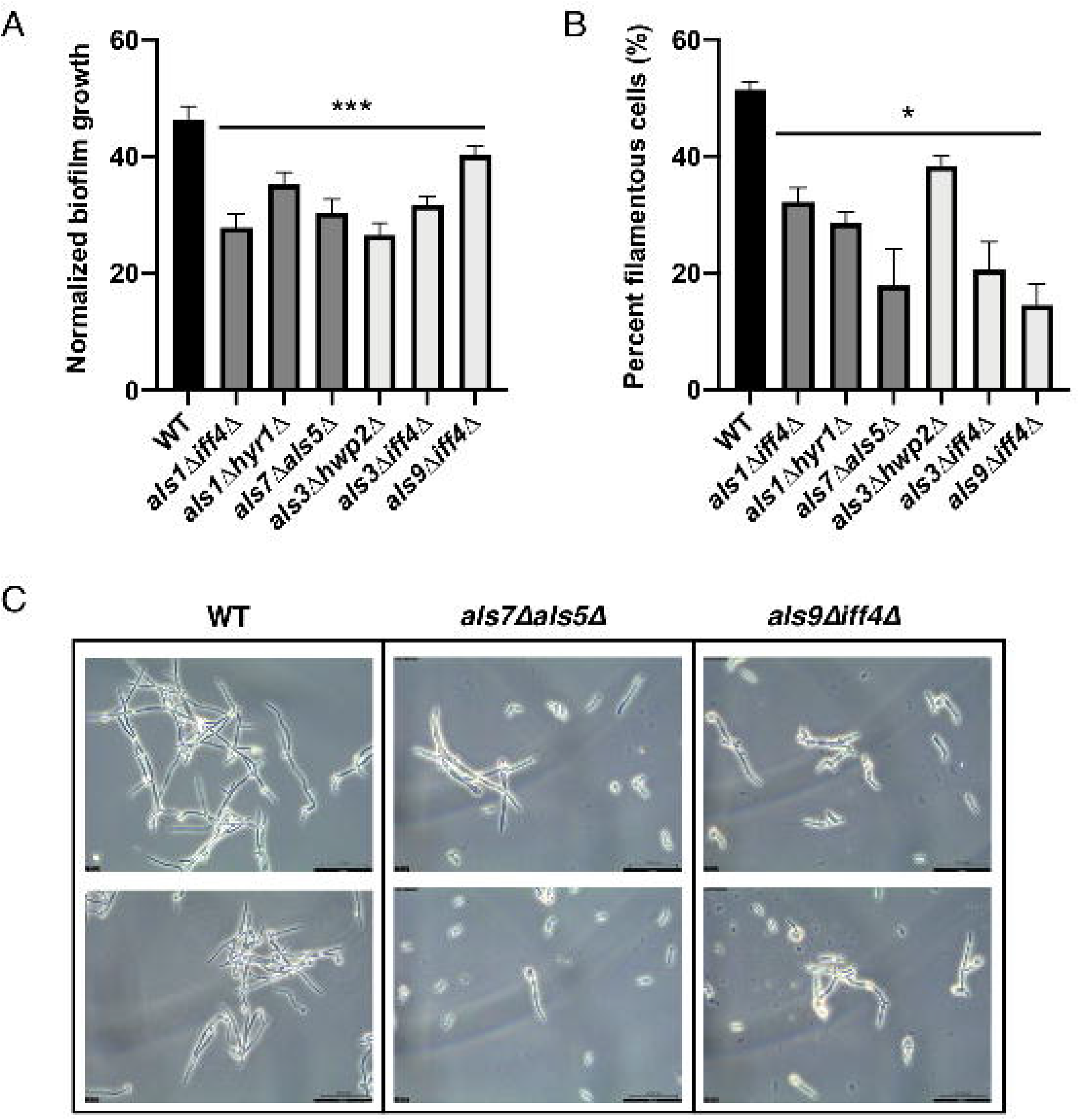
Adhesin mutant strains are deficient in filamentation and biofilm growth. The three strains with lowest virulence (*als1Δiff4*, *hyr1Δals5Δ*, *als7Δals5Δ*) and three with highest virulence (*als3Δhwp2Δ*, *als3Δiff4Δ*, *als9Δiff4Δ*), based on the high-throughput *C. elegans* liquid media screen, have reduced ability to undergo morphogenesis or form biofilms, regardless of virulence phenotype. **(A)** Lowest virulence strains (*als1Δiff4*, *hyr1Δals5Δ*, *als7Δals5Δ*; dark grey) and highest virulence strains (*als3Δhwp2Δ*, *als3Δiff4Δ*, *als9Δiff4Δ*; light grey), all have reduced ability to form biofilms compared to a wild-type strain (ANOVA, *P* < 0.0005 (***)). Biofilm growth was quantified by an XTT metabolic readout, measured at OD490 and normalized to planktonic growth. **(B)** Lowest virulence strains (*als1Δiff4*, *hyr1Δals5Δ*, *als7Δals5Δ*; dark grey) and highest virulence strains (*als3Δhwp2Δ*, *als3Δiff4Δ*, *als9Δiff4Δ*; light grey), all have reduced ability to form filamentous cells (ANOVA, *P* < 0.05 (*)). *C. albicans* cells were grown in media containing 10% fetal bovine serum (FBS) at 37°C to induce filamentation, and cells were counted using brightfield microscopy to determine the percentage of filamentous cells in the population. **(C)** Examples of reduced filamentation in a lower virulence (*als7Δals5Δ*) and higher virulence (*als9Δiff4Δ*) mutant strain. *C. albicans* cells were grown in media containing 10% FBS at 37°C to induce filamentation, and cells were counted using brightfield microscopy. Two representative microscopy images are shown for each strain at 40X magnification, scale bar is 50 μm.

Selected mutants used in follow up biofilm assays were also used in filamentation assays to assess the filamentation capabilities of adhesin mutants with variable virulence profiles. For this assay, wild-type and mutant strains were cultured in media with or without 10% serum, to induce filamentation in *C. albicans* cells. Each strain was imaged under phase microscopy and the percentage of filamentous cells (as a fraction of total cells) was calculated. Similar to what was observed with the biofilm growth assay, each of the adhesin mutant strains were found to have impaired ability to undergo morphogenesis and grow as filamentous cells, compared with the wild-type strain (**Figure 5b**; *P* < 0.05); although each of these mutants retains the ability to filament, they formed fewer filamentous cells compared with a wild-type strain. Representative microscopy of a low-virulence (*als7Δals5Δ*) and high-virulence (*als9Δiff4Δ*) strain indicate that both low- and high-virulence adhesin mutants are impaired in filamentous growth compared with a wild-type strain, when grown in the presence of serum (**Figure 5c**). The role of adhesins in mediating *C. albicans* filamentation has been less well-studied compared with biofilm formation. This data suggests that deleting adhesin factors results in reduced ability to form filamentous cells, but similar to biofilm formation, is not correlated with virulence in our *C. elegans* liquid or agar infection assays.

### Genetic interaction analysis identifies significant negative interactions amongst C. albicans mutants

In addition to identifying the most virulent and avirulent adhesin mutants, we further wanted to characterize any potential genetic interactions between adhesin genes. Genetic interaction analysis would allow us to identify significant positive and negative genetic interactions, which point towards important synergies between these adhesin factors. Therefore, we performed genetic interaction analysis on our *C. albicans-C. elegans* virulence datasets, which compares double mutant deletion strain fitness to that of their single mutant counterparts (*i.e. aΔbΔ* fitness compared to *aΔ* fitness and *bΔ* fitness). We used the commonly-employed multiplicative model of genetic interactions (Boone *et al.* 2007; Baryshnikova *et al.* 2013; Halder *et al.* 2019), which predicts that the fitness of a double mutant will be the product of the fitness of each single mutant counterpart. Double mutant strains found to be less fit than predicted are said to have a negative genetic interaction, and those with higher fitness have a positive genetic interaction. For our assay, negative genetic interactions are particularly interesting, as it suggests deleting two adhesins in combination may lead to significantly reduced virulence — more than would be expected by deletion of either single adhesin on its own.

We used a simple program to assess genetic interactions and compare each double adhesin deletion mutant to its single mutant counterparts. First, for each replicate of the virulence screen, we normalized each mutant to the wild-type strain in order to obtain a relative fitness score. We then used these relative fitness measurements to calculate a predicted interaction score for each double mutant, based on the multiplicative model. Each predicted score was compared to the double mutant scores for both reciprocal pairs in each of the six replicate assays, and *t*-tests were used to compare the experimental values and the predicted values and determine whether we would reject the null that there is no difference between the two (**Figure 6A**). We found two double mutants to have a significant negative genetic interaction: *als5Δeap1Δ* and *als5Δhwp2Δ* (**Figure 6A, 6B**), *P* < 0.05. This indicates that deletions of *als5Δ* and *eap1Δ*, or *als5Δ* and Δ*hwp2Δ* together, renders *C. albicans* significantly less virulent than deletion of *als5Δ*, *eap1Δ*, or Δ*hwp2Δ* or their own, and that these factors act synergistically to promote fungal virulence in this *C. elegans* infection model.

**Figure 6.**
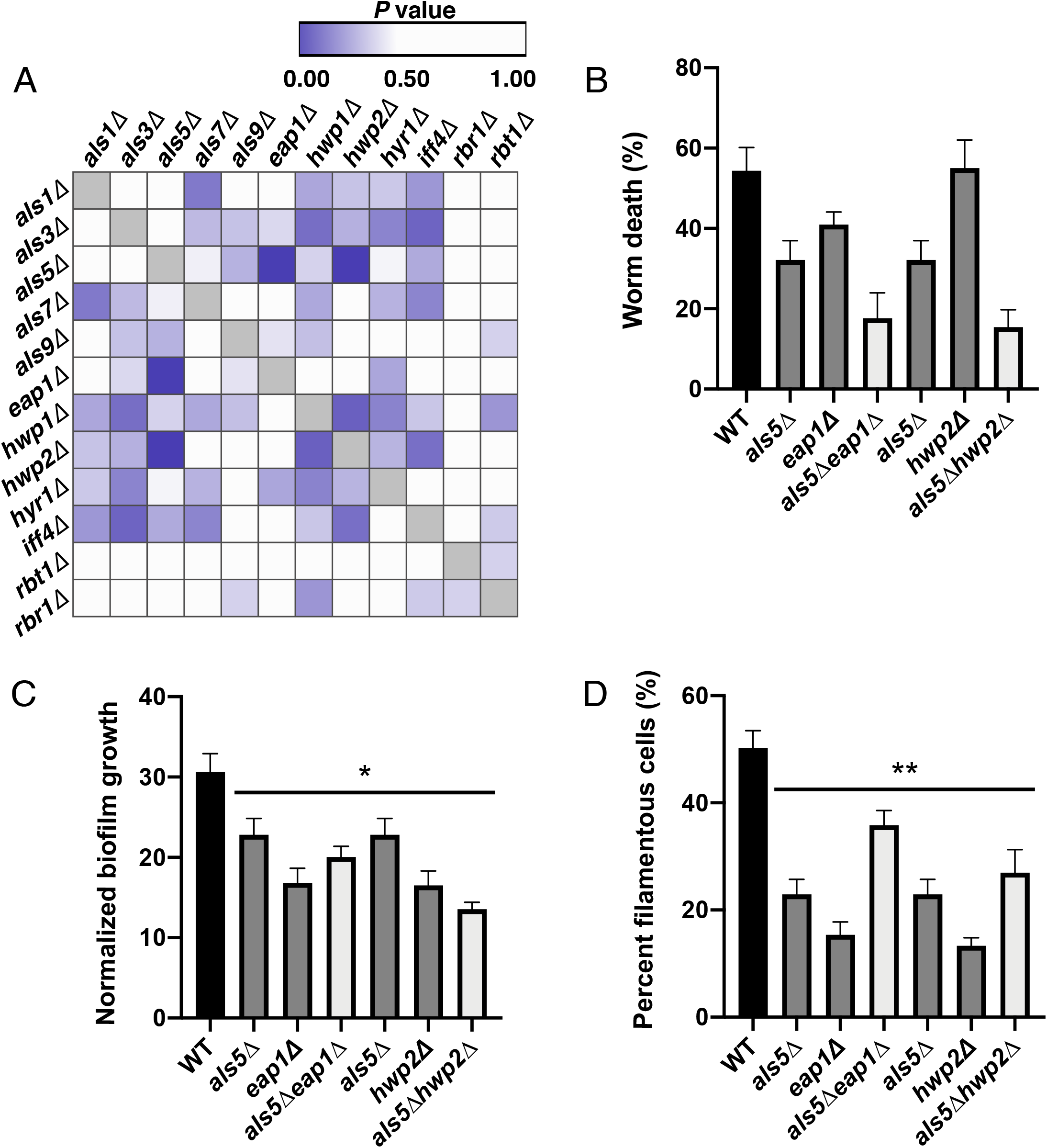
Genetic interaction analysis of adhesin mutant strains. Genetic interaction analysis was performed for all adhesin mutant strains to identify and characterize genetic interactions. **(A)** Summary of genetic interactions between each of the double mutant adhesin strains. A multiplicative model of genetic interactions was used to identify interactions where the double mutant strain virulence (fraction of dead worms) deviated from the predicted virulence based on the product of the two corresponding single mutants. The heatmap shows *P* values for each double mutant strain (based on actual vs. predicted virulence measure) across the six replicate experimental infection assays. Darker blue/purple represents lower *P* values, and white represents higher values. Two mutants (als5Δeap1Δ and als5Δhwp2) were identified as significant interactions with *P* < 0.05. The heat map was generated using Morpheus matrix visualization and analysis software from the Broad Institute (https://software.broadinstitute.org/morpheus). **(B)** The virulence (percent worm death) of the two genetic interaction mutants (light grey), compared with their single mutant constituent mutants (dark grey) as well as the wild-type strain, based on the high-throughput *C. elegans* liquid media screen. Graph depicts the percent of worm death. **(C)** Each of the genetic interaction double mutants and their single mutant constituent strains have reduced ability to form biofilms compared to a wild-type strain (ANOVA, *P* < 0.05 (*)). Biofilm growth was quantified by an XTT metabolic readout, measured at OD490 and normalized to planktonic growth. **(D)** Each of the genetic interaction double mutants and their single mutant constituent strains have reduced ability to form filamentous cells (ANOVA, *P* < 0.005 (**)). *C. albicans* cells were grown in media containing 10% FBS at 37°C to induce filamentation, and cells were counted using brightfield microscopy to determine the percentage of filamentous cells in the population.

Finally, we assessed whether these two genetic interaction mutants (*als5Δeap1Δ* and *als5Δhwp2Δ*) were impaired in filamentation and biofilm formation, compared with their counterpart single mutant strains. We performed a biofilm growth assays with each of these mutants, and found that while the single mutant strains (*als5Δ*, *eap1Δ*, and Δ*hwp2Δ*) as well as the double mutant strains (*als5Δeap1Δ* and *als5Δhwp2Δ*) were all impaired in biofilm formation, compared to the wild-type strain (**Figure 6C**, *P* < 0.05), the double mutants were not further deficient in biofilm growth compared to their single mutant counterpart strains. Similarly, all single and double mutants strains were significantly impaired in filamentation compared to the wild-type strain (**Figure 6D**, *P* < 0.005), yet the double mutants were not further deficient in filamentation growth compared to their single mutant counterpart strains, and in fact filamented more robustly than their single mutant counterparts. This suggests an uncoupling between filamentation, biofilm formation, and virulence, and/or further indicates how specific environmental conditions may influence the requirement for different adhesin proteins.

## Discussion

Here, we present a systematic analysis of the role of adhesin factors, singly and in combinations, in *C. albicans* virulence. Our initial screening of 144 *C. albicans* strains exploited *C. elegans* as a model host uniquely suited for such high-throughput virulence analysis, and revealed single- and double-genetic mutant strains that retained high levels of virulence (comparable to wild type), and strains with significantly reduced levels of virulence. This screening platform was further able to identify genetic interactions between adhesin genes, and specific pairs of mutants that are significantly less virulent that would be expected based on the virulence of their single mutant constituent strains. This highlights the value in assessing virulence in higher-order genetic mutants, such as these double mutant libraries. In addition to screening for virulence, we followed up on strains with the highest and lowest levels of virulence. We found that *in vitro* measures of pathogenicity traits, including *C. albicans* filamentation, and biofilm formation were impaired in adhesin mutant strains regardless of their virulence patterns in *C. elegans*, suggesting that under the conditions tested, filamentation and biofilm growth were uncoupled from virulence, which has been previously suggested (Noble *et al.* 2010). Further, we found that mutants with less virulence in the *C. elegans* agar model were less well able to colonize these worms, compared with the more virulent strains, and that *C. elegans* survival upon *C. albicans* infection varied under different infection conditions (*i.e.* solid agar vs. liquid killing assays). Together, these data suggest a complex role of adhesin factors in mediating fungal virulence, and that specific environmental conditions have a critical role in influencing the requirement for different adhesin factors.

Host-pathogen interactions are complex and dynamic. In this manuscript, we demonstrated that *C. albicans* likely utilizes different virulence mechanisms in liquid- and agar-based assays. First, colonization on agar is higher than in liquid pathogenesis conditions (**Figure S2**). Second, mutants with higher colonization (*als1Δiff4Δ*, *als1Δhyr1Δ*, *als7Δals5Δ*) are more virulent in an agar-based assay compared to mutants with lower colonization (*als3Δhwp2Δ*, *als3Δiff4Δ*, *als9Δiff4Δ*). At the same time, these higher colonization mutants have an approximately 5-fold decrease in their ability to kill worms in the liquid-based assay. Interestingly, the same result was observed in the well-studied *C. elegans* – *P. aeruginosa* pathosystem, where multiple virulence models have been developed. For example, agar-based Slow Killing is characterized by high colonization of the host and the requirement for bacterial quorum sensing pathways for full virulence, with *lasR* and *gacA* mutants being low colonizers with attenuated virulence (Tan *et al.* 1999; Feinbaum *et al.* 2012). In contrast, Liquid Killing is characterized by low colonization, wild-type virulence of quorum-sensing mutants, and a requirement for the siderophore pyoverdine for full virulence (Kirienko *et al.* 2013, 2015; Kang *et al.* 2018). In this assay, multiple *pvd* mutants are attenuated, but no difference in colonization is observed between bacteria with low- and wild-type virulence (Kirienko *et al.* 2013). This suggests that there may be a wide-spread correlation between colonization and virulence in agar assays but that correlation will be absent in liquid-based assays, where colonization is reduced.

The clear importance of different infection models and the broader pathogen environment on influencing fungal virulence, highlights the need to assess pathogenicity in different contexts. While mammalian models such as the mouse intravenous tail vein infection model for systemic candidiasis remain a gold standard for fungal virulence assays (Segal and Frenkel 2018), the simplicity of the *C. elegans* model and its tractability for high-throughput manipulation allowed us to rapidly screen a large library of adhesin mutants, and identify key regulators of virulence. Indeed, a breadth of previous research has exploited this model to study fungal virulence (Pukkila-Worley *et al.* 2009), the role of the antifungal host immune response (Pukkila-Worley *et al.* 2011, 2014), and antifungal drug efficacy (Breger *et al.* 2007; Okoli *et al.* 2009; Ewbank and Zugasti 2011) in *C. albicans*, and other fungal pathogens (Mylonakis *et al.* 2002a; Scorzoni *et al.* 2013). Other infection models that have been valuable for the study of *C. albicans* virulence include a *Drosophila* infection model (Alarco *et al.* 2004; Chamilos *et al.* 2006; Glittenberg *et al.* 2011; Wurster *et al.* 2019), *Galleria mellonella* moth larvae model (Fuchs *et al.* 2010; Frenkel *et al.* 2016). Very recently, *Manduca sexta* caterpillars have been developed as a novel host model for the study of fungal virulence and drug efficacy (Lyons *et al.* 2020), which have several advantages over other non-mammalian models, including their ability to be maintained at 37°C and ability to assess fungal burden throughout the course of infection via the caterpillar’s hemolymph of feces. Other recent work has identified a novel virulence phenotype to assess *Candida* pathogenesis in the *C. elegans* host model; this study found that in addition to causing host lethality, fungal pathogens such as *C. albicans* also reduce *C. elegans* fitness by delaying reproduction (Feistel *et al.* 2019). This has longer-term implications for overall *C. elegans* population growth, adds an important new layer to our understanding of this host-pathogen interaction, and may provide a more complete picture of virulence when studying fungal mutant strains, such as the adhesin mutants described here.

One of the unique capabilities of a high-throughput host-pathogen interaction model, is our ability to perform complex genetic interaction analysis. Genetic interaction analysis typically requires the analysis of single- and double-genetic mutant strains to compare double mutants to their single mutant counterparts, and often requires the analysis of numerous strains in order to identify significant interactions. While this work represents one of the first genetic interaction screens to monitor *C. albicans* virulence using an *in vivo* infection model, numerous other studies have probed the genetic interactions networks mediating pathogenesis traits in fungal pathogens (Bharucha *et al.* 2011; Diezmann *et al.* 2012; Usher *et al.* 2015; Glazier *et al.* 2017, 2018; Shapiro *et al.* 2018a; Glazier and Krysan 2020), as well as numerous bacterial pathogens (Joshi *et al.* 2006; van Opijnen *et al.* 2009; Côté *et al.* 2016; Skwark *et al.* 2017), and parasites (Fang *et al.* 2018). Genetic interaction analysis in microbial pathogens has similarly been used as a means to probe complex genetic networks mediating host-pathogen interactions (O’Connor *et al.* 2012; Urbanus *et al.* 2016; Lee *et al.* 2019).

In addition to using genetic interaction analysis as a means to understand genetic networks, genetic interaction analysis can also be exploited as a means to uncover novel pairs of cellular targets for combination antimicrobial therapeutics. In particular, synthetic lethal interactions, where mutation of two genes in combination is lethal to the cell while deletion of either gene on its own remains viable, can be used to identify targets for combination drug therapies (Cokol *et al.* 2011). Such genetic interaction-based approaches have been well validated for combination therapies for cancer (Han *et al.* 2017), as well as for antimicrobial therapeutics (Cheng *et al.* 2014; Pasquina *et al.* 2016; Usher and Haynes 2019). While our work does not identify lethal genetic combinations, we have identified genetic combinations that significantly impair virulence, more than would be expected by mutating the constituent single genes. Targeting virulence regulators, which impair a pathogen’s ability to cause infection without altering its overall fitness, is gaining momentum as a potentially effective strategy for antimicrobial therapy (Cegelski *et al.* 2008; Maura *et al.* 2016; Dickey *et al.* 2017). Such ‘antivirulence’ agents have been identified that inhibit pathogenicity traits such as morphogenesis and biofilm formation in *C. albicans* (Toenjes *et al.* 2005; Fazly *et al.* 2013; Vila *et al.* 2017; Romo *et al.* 2017; Garcia *et al.* 2018). Our work lends an understanding to new combinations of adhesins that significantly impair fungal virulence, suggesting new putative targets for combination antivirulence therapeutics.

## Materials and Methods

### C. elegans-C. albicans liquid infection assay and screening

The *C. elegans* - *C. albicans* infection assay was performed with slight modifications to the previous method (Kirienko *et al.* 2014). In brief, *C. elegans glp-4(bn2)* mutants were reared on NGM media seeded with concentrated *E. coli* OP50 media until they reached young adult stage (1 day at room temperature and then shifted to 25 °C for 2 days). *C. albicans* strains were cultured overnight in 96-well deep-well plates containing 300 μL YPD and incubated at 37 °C. OD_600_ of plate was read and cultures were diluted with S Basal to normalize *C. albicans* density. 384-well assay plates were set, with final media composition of 10% BHI, *C. albicans* at OD_600_=0.03, 3.5 μg/mL cholesterol, and S Basal (to bring the final volume to 50 μl/well). Using a COPAS FlowPilot (a large-bore fluorescence-activated worm sorter, analogous to a FACS machine), and LP (large particle) Sampler (Union Biometrica, MA), 25 *C. elegans* nematode worms were added to each well. Plates were covered with breathable membranes and incubated for 72 hours at 25 °C. OD_600_ was read for each plate. Plates were washed 5 times with S Basal then liquid was aspirated using an EL 406 Plate washer / Liquid dispenser (BioTek, VT). 50 μL of 0.01% tween was added to each well to limit worms adhering to pipette tips; worms were then transferred to new 384-well plates to minimize background. Plates were washed 5 times with S Basal to remove transferred pathogens, and then liquid was aspirated. 50 μL of SytoxOrange stain (0.2 μL Sytox /1mL S Basal) was loaded into each well. Plates were incubated 12-16 hours in a dark place at room temperature. Plates were washed 5 times with S Basal. Plates were imaged with both bright field and red channel RFP using a Cytation 5 plate reader (BioTek, VT). Images were analyzed for worm survival using CellProfiler, as previously described (Anderson *et al.* 2018). Statistical analysis was performed on all data to ensure significance. *C. elegans* liquid infection assay data was initially checked using the coefficient of variation (CV) and outliers were removed to obtain CV scores of 0.5 or less in as many strains as possible. The remaining strains had CV scores of less than 1. The selected mutants in all datasets were subjected to one-way ANOVA tests to compare each strain to the wild-type.

### Genetic interaction analysis

Each single mutant tested in this research was assigned a fitness score based on the results of the *C. elegans* infection assay. The fitness score of the wild-type was normalized to 1, and the results from all other strains were divided by the wild-type results to achieve these relative fitness scores. The fitness scores of each single parent strain were multiplied by one another and the product became the predicted fitness score of the double mutants. Genetic interaction analysis was based on deviation from predicted fitness score. If the double mutant’s actual score was significantly different from its predicted score it was considered a “hit”. Genetic interaction analysis was done using R. First, the program bound all the replicate results together, labelled each single mutant and created a single dataset with the library’s unique combinations. A two-sided t-test was used to compare the predicted and actual fitness scores. A positive interaction had an adjusted *P* value < 0.05 and a deviation that is greater than the predicted score. A negative interaction had an adjusted *P* value < 0.05 and a deviation that is lower than the predicted score. The R program used for this GI analysis can be found here: https://github.com/kieran11/wormdata

### Colonization assay

To assess *C. elegans* colonization by *C. albicans* following agar- or liquid-based infection, worms were infected as described above. After 24 hours, worms were transferred onto a clean plate and allowed to crawl away from *C. albicans*. Then they were transferred via pick to a 1.5 mL microcentrifuge tube containing S Basal with 1% levamisole, and incubated 10 minutes till all worms were paralyzed. Worms were washed 5-6 times in S Basal containing 0.01% Triton X-100 to remove residual fungi. After the final wash, worms were resuspended in 200 μL of 0.1% Triton-X 100 in S Basal and vortexed for 1 min using zirconium beads (Thermofisher Scientific, NC0442292). The resulting lysate was serially diluted 5-fold and plated onto YPD agar plate to quantify the number of colony forming units (CFU) per unit volume. This value was then used to derive the average number of CFUs per worm in each sample. The assay was performed using three biological replicates, each of them consisting of three technical replicates, with 15 worms per technical replicate.

### C. elegans-C. albicans agar-based killing assay

Kim et al. recently outlined an agar killing assay as a nematode infection model to study fungal pathogenesis (Kim *et al.* 2020). Briefly, *C. elegans glp-4* eggs were extracted with worm bleach and resuspended in S Basal to establish a fresh stock of worms. The *C. elegans* worms were then counted on an NGM plate under a stereomicroscope and 5000-6000 worms dropped onto two superfood agar plates, which were left overnight at room temperature then moved to 25°C for an additional two nights. *C. albicans* strains were incubated overnight in 1 mL of YPD at 37°C and 250 rpm and then 70μL of culture was spread onto a BHI-Kan plate, which were then incubated for 16 hours at 30°C. Approximately 6 mL of S Basal was dispensed onto each *C. elegans* superfood agar plate to dislodge the worms. The worms were then washed 3 times with S Basal before being made into a solution of 2 worms per μL of S Basal. 45-55 worms were transferred onto each *C. albicans*-inoculated BHI-Kan plate, which were then left at room temperature for 1 hour before being stored overnight at 25°C. Dead worms were counted and removed with a platinum wire worm pick from the plates under a stereomicroscope daily until all worms were dead.

### qPCR

For quantitative, real-time reverse-transcription PCR (qPCR), *C. albicans* were prepared as for agar- or liquid-based assays. Next, *C. albicans* was either scraped off the plate and resuspended in S Basal or centrifuged down (for agar or liquid assays, respectively). RNA was extracted from an ~100 μL fungal pellet. RNA was purified by vortexing the fungal sample with glass beads and then subsequent Trizol extraction with BCP as a phase-separating agent, as previously described (Kirienko *et al.* 2018). Reverse transcription was performed using an AzuraQuant cDNA Synthesis Kit (Azura). qPCR was conducted in a CFX-96 real-time thermocycler (Bio-Rad) using SYBR green AzuraQuant Fast Green Fastmix (Azura). Fold-changes were calculated using a ΔΔCt method with actin as a housekeeping gene. Cycling parameters and primer sequences are available upon request. For each experiment, at least three biological replicates were performed.

### Filamentation assay and analysis

*C. albicans* strains were incubated overnight in 5 mL YPD at 37°C and 350 rpm. OD_600_ of each culture was taken. Cultures were diluted with fresh YPD media to normalize to the same OD_600_ value. *C. albicans* was subcultured into 5 mL YPD with 10% fetal bovine serum (FBS). Subcultures were incubated at 37°C and 350 rpm for 4 hours. 10μL of each culture was pipetted onto a microscope slide and visualized under phase microscopy at 40X magnification using a Leica DM 2000 LED microscope. Images of each strain were taken with a Leica ICC50 W camera. Images were analyzed using the ImageJ program and MicrobeJ plugin. Analysis involved using ImageJ to count the number of yeast cells and filamentous cells in each image as well as measure the length of each filamentous cell in each image.

### Biofilm assay

*C. albicans* strains were incubated overnight in 5 mL YPD at 37°C and 350 rpm. OD_600_ of each culture was taken. Cultures were diluted with BHI+Kan media to normalize to the same OD_600_ value. Strains were subcultured into flat bottomed 96-well polystyrene plates containing 100 μL of BHI+Kan and 100 μL of culture per well. 200 μL of YPD per well in one row acts as a negative control. 96-well plates were wrapped in tinfoil and incubated at 37°C for 72 hours. 120 μL of planktonic cells from each well was transferred to new 96-well plates, then read in a plate reader at OD_600_. Plates were washed twice with 200μL PBS and left upside down to dry in a fume hood. 1 hour later or once dry, 90 μL of XTT (1 mg/mL PBS) and 10ul of PMS (0.32 mg/mL H_2_O) was added to each well. Plates were wrapped in tinfoil and incubated for 2 hours at 30°C. Plates were read at 490nm using a Tecan infinite M nano spectrophotometer. The XTT biofilm growth values were normalized to planktonic cell growth.

### Data and reagent availability

*C. albicans - C. elegans* virulence screening data is available in **Supplemental Table S1**. The R program used for this GI analysis can be found here: https://github.com/kieran11/wormdata. All *C. albicans* strains used as part of this research will be made available upon request.

## Acknowledgements

Funding for this research was provided by the Canadian Institutes for Health Research, the Natural Sciences and Engineering Research Council to R.S., a Mitacs Globalink Research Award to S.R., a University of Guelph, College of Biological Science CBS Undergraduate Summer Research Assistantship Award to G.K, and a Welch Foundation and a John S. Dunn Foundation Award to N.V.K. We would like to thank the members of the Shapiro lab and the Kirienko lab for their hard work and assistance. Thank you to Kieran Shah for his help writing the genetic interaction analysis script and to Anastasia Baryshnikova for helpful discussions.

